# Developmental Elimination of Electrical Synapses by UNC-51/UNC-76-Mediated Vesicular Transport

**DOI:** 10.64898/2026.03.04.709731

**Authors:** Hao Huang, Yuzhi Yang, Shujie Qiu, Yuanhao Xu, Yuli Jian, Zihao Zhao, Dong Yan, Lingfeng Meng

## Abstract

Neural circuit maturation requires precise elimination of transient synaptic connections, but how electrical synapses are eliminated remains unknown. Using *C. elegans* PLM mechanosensory neurons, we found that gap junctions form transiently during early development and are subsequently eliminated as the circuit matures. These transient electrical synapses drive high-frequency calcium oscillations, and optogenetic ablation of early gap junctions abolishes calcium dynamics and disrupts chemical synapse formation, demonstrating their critical developmental function. Through forward genetic screening, we identified the conserved kinase UNC-51/ULK as essential for elimination. UNC-51 phosphorylates the kinesin adaptor UNC-76/FEZ to initiate RAB-10-dependent retrograde trafficking, progressively depleting innexin proteins from the synaptic site. Disrupting this pathway prevents elimination, causing persistent electrical synapses and neuronal hyperactivity. These findings establish a conserved framework for developmental elimination of functionally essential transient electrical connectivity.

## Introduction

Neural circuit maturation relies on the precise elimination of transient synaptic connections to achieve functional specialization^1^. While mechanisms governing the pruning of chemical synapses are well characterized—including activity-dependent processes such as complement-mediated tagging and microglial phagocytosis^2–8^—how electrical synapses are eliminated during development remains fundamentally unknown.

Electrical synapses, formed by connexin/innexin gap junction channels that create intercellular pores permeable to ions, calcium, and second messengers^9,10^, enable not only rapid electrical signal transmission but also direct intercellular chemical signaling. These channels exhibit remarkably rapid turnover (1-5 hours)^11^—among the fastest in the proteome—due to continuous cycles of membrane insertion and removal^12,13^. This dynamic trafficking enables precise regulation of electrical coupling strength, which is directly determined by the density of gap junctions at intercellular junctions^14,15^. While activity-dependent transcriptional mechanisms can modulate expression levels, the trafficking machinery itself provides an additional layer of control over synaptic connectivity^16^. However, whether organisms actively exploit this trafficking machinery to programmatically adjust intercellular communication during development remains largely unexplored.

Transient electrical synapses are a conserved and widespread feature of developing neural circuits, representing the earliest functional connections to form across systems from invertebrates to mammals^17–20^. During this early developmental window, they serve remarkably diverse functions: coordinating left-right asymmetry in *C. elegans* AWC neurons^21^, driving spontaneous activity waves for circuit refinement in mammalian retina^22^; and synchronizing spinal interneuron networks to drive the earliest embryonic movements in zebrafish^23^. Remarkably, these early electrical connections are not merely active during this window—they are functionally required for the formation of the chemical synapses that will eventually replace them: disrupting gap junction expression in leech neurons prevents chemical synaptogenesis^24^, while blocking electrical coupling between sister pyramidal cells in mouse cortex impairs their later chemical connectivity^25^. Although gap junctions are known to transmit calcium and second messengers, whether these signals causally instruct chemical synapse development—and how these transiently essential connections are themselves subsequently eliminated—remain entirely unknown.

Here, using *C. elegans* PLM mechanosensory neurons, we address both questions. We show that transient electrical synapses coordinate high-frequency calcium oscillations during early larval development, and that loss of these synapses abolishes calcium dynamics and disrupts chemical synapse formation—directly linking electrical coupling, calcium signaling, and chemical synaptogenesis within a single system. Through forward genetic screening, we identify the conserved kinase UNC-51/ULK as the central regulator of electrical synapse elimination, acting through a phosphorylation-dependent trafficking mechanism that drives directional removal of innexin proteins from synapses. These findings define the developmental function of transient electrical synapses in instructing circuit assembly, and uncover a conserved kinase-driven trafficking mechanism that eliminates them once this function is fulfilled.

## Results

### Electrical synapses undergo developmental elimination in mechanosensory neurons

To investigate electrical synapse regulation during development, we focused on PLM mechanosensory neurons in *C. elegans*, which form electrical connections with their partner neurons LUA, PVC and PVR ^26,27^. Electrical synapses were visualized using the stable GFP::UNC-9 transgenic line (*yadIs12*) since UNC-9 is a major gap junction component in PLM neurons and GFP::UNC-9 puncta have been validated to mark electrical synapses^28^. Quantification of electrical synapse numbers along PLM neurites across L1 (first larval stage) to L4 (fourth larval stage) revealed that L1 displayed approximately 8 electrical synapses (median=8) distributed along the entire neurite. As development progressed, electrical synapse numbers significantly decreased by L2 (median=5) and stabilized at reduced levels during L3 and L4 (median=4 for both stages) (Fig. 1A,C). Notably, the most pronounced decline occurred between L1 and L2 (Fig. 1C), demonstrating that the majority of electrical synapse elimination takes place by L2.

**Figure 1.**
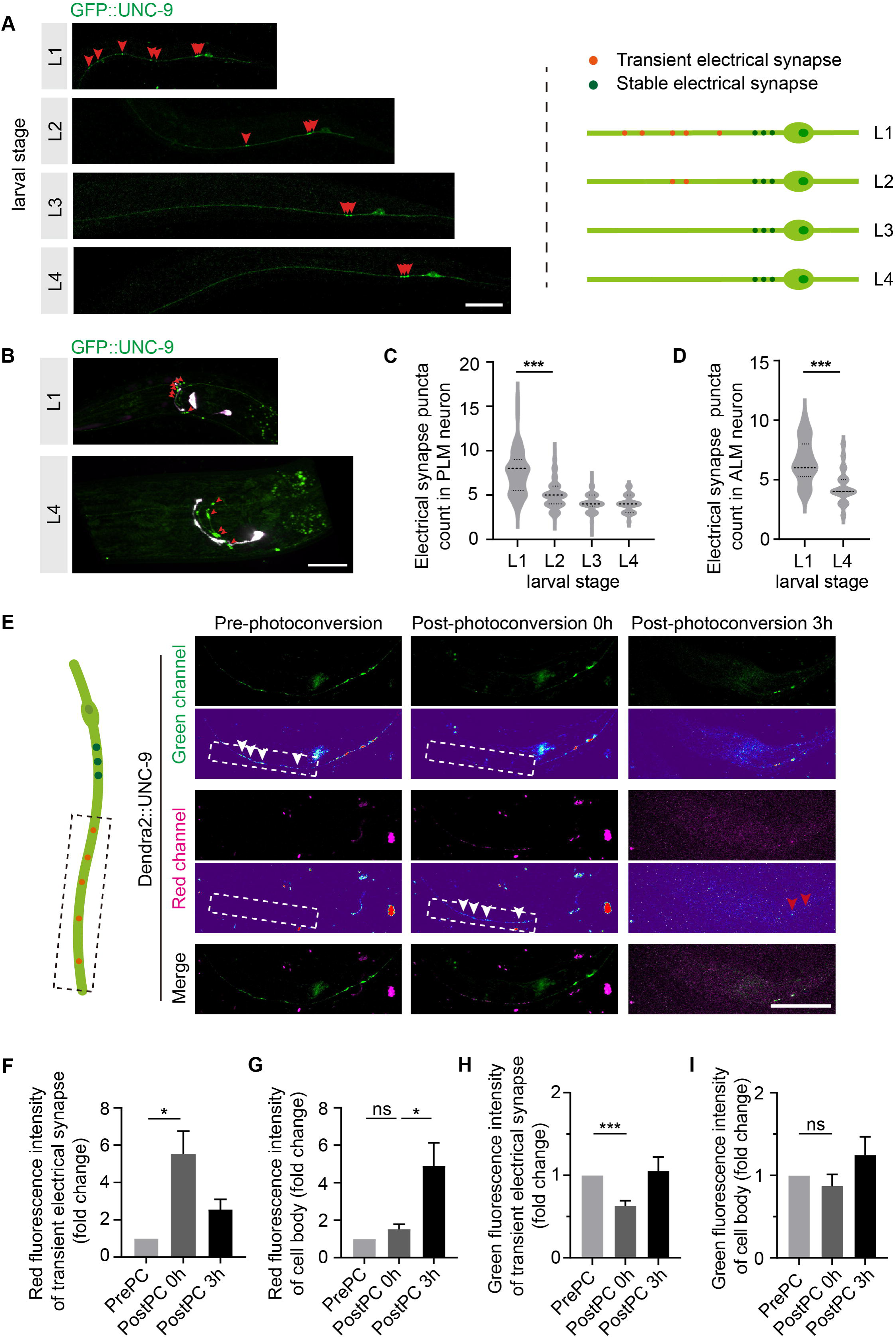
Transient electrical synapses are developmentally eliminated in mechanosensory neurons. (A) Representative fluorescence images of GFP::UNC-9-labeled electrical synapses in PLM neurons across larval development (L1, L2, L3, and L4) in *yadIs12* animals. Red arrowheads point to electrical synapses in the left panel, while the right panel presents a schematic of the L1–L4 PLM electrical synapse elimination. (B) Representative fluorescence images of GFP::UNC-9-labeled electrical synapses in ALM neurons at L1 and L4 in *yadIs12* animals. Red arrowheads point to electrical synapses. (C) Quantification of GFP::UNC-9-labeled electrical synapses in PLM neurons across larval development in *yadIs12* animals. n > 100 per stage. The dashed lines represent the quartiles and median of the data distribution. (D) Quantification of GFP::UNC-9-labeled electrical synapses in ALM neurons at L1 and L4 in *yadIs12* animals. n = 20 (L1), n = 24 (L4). (E) Representative images of photoconvertible Dendra2::UNC-9 on PLM before photoconversion, immediately post-photoconversion, and 3 hours post-photoconversion, shown in red and green channel, with corresponding heat map images indicating signal intensity. White arrowheads point to Dendra2 labeled UNC-9 puncta on the PLM neurites, and red arrowheads point to Dendra2 labeled UNC-9 in the cell body. (F-I) Quantification of red and green Dendra2::UNC-9 fluorescence signal in transient electrical synapse area and cell body at pre-photoconversion, immediately post-photoconversion, and 3 hours post-photoconversion. n = 9 per condition. For panels with quantification, error bars represent the mean ± SEM. For confocal images, scale bar: 20 μm. Data in (D) was analyzed by Student’s t-test. (C), (F), (G), (H) and (I) were analyzed by one-way ANOVA with Tukey’s multiple comparisons test. ns: not significant, *p < 0.05, ***p < 0.001.

For confirmation that the observed elimination pattern reflects endogenous protein dynamics rather than transgenic artifacts, CRISPR-mediated endogenous tagging of UNC-9 with mNeonGreen was performed. Endogenously tagged UNC-9::mNeonGreen in PLM neurons showed a similar developmental decline from L1 to L4 (Fig. S1A), validating our findings from the transgenic line.

To determine whether this phenomenon extends beyond PLM neurons, we examined ALM mechanosensory neurons, which form electrical connections with AVD interneurons in the nerve ring through UNC-9 and UNC-7 innexins^29,30^. Using the *yadIs12* transgenic reporter, we quantified UNC-9 puncta in ALM across development. At L1, ALM neurons displayed approximately seven puncta (median = 7), which significantly decreased to approximately four puncta (median=4) by L4 (Fig. 1B, D). Together, these results demonstrate that electrical synapse elimination is not restricted to PLM neurons, but represents a conserved developmental process in *C. elegans* mechanosensory neurons.

Given the rapid elimination in electrical synapse at early stages, we performed time-lapse imaging of HaloTag::UNC-9 in L1 PLM neurons to examine its dynamic behavior. This analysis revealed that transient electrical synapses undergo continuous assembly and disassembly, with formation and dissolution rates occurring at comparable levels along the neurite (Fig. S1B,C), resembling the behavior of stable electrical synapses in young adult animals^28^. These observations suggested that transient electrical synapses also exhibit highly dynamic properties. To further validate the dynamic behavior of electrical synapses and quantify protein turnover rates, photoactivation experiments were performed using photoconvertible Dendra2::UNC-9 in L1 PLM neurons. Selective photoactivation of Dendra2::UNC-9 at transient electrical synapses induced a conversion from green to red fluorescence (boxed region, Fig. 1E), and the red fluorescent signal was monitored at 0 and 3 hours post-activation. Following photoactivation, green fluorescence at the targeted region decreased by half, whereas red fluorescence increased approximately six-fold compared with the pre-activation baseline, indicating efficient photoconversion (Fig. 1F,H). The fluorescence signal in the soma remained unchanged (Fig. 1G, I), demonstrating specificity of the photoactivation. Time-course imaging over 3 hours revealed a progressive decline in red fluorescence at electrical synapses, with intensity decreasing by half while simultaneously increasing in the soma (Fig. 1F,G), indicating active protein retrieval with a half-life of approximately 3 hours. This turnover rate is comparable to that of stable electrical synapses measured at L4, demonstrating that transient synapses undergo similarly rapid turnover.

### Transient electrical synapses coordinate calcium dynamics to regulate chemical synapse development

To determine the functional role of transient electrical synapses, we next examined spontaneous calcium activity in PLM neurons at L1 (electrical synapses present) and L4 (electrical synapses eliminated) using GCaMP6. Both L1 and L4 animals exhibited typical PLM calcium responses to mechanical stimulation (Fig. 2C), confirming that sensory function is intact across developmental stages. However, wavelet transform analysis of calcium dynamics revealed that L1 neurons exhibited significantly higher power in high-frequency components compared to L4 (Fig. 2A,E), indicating that electrical synapses enhance high-frequency calcium oscillations during early development.

**Figure 2.**
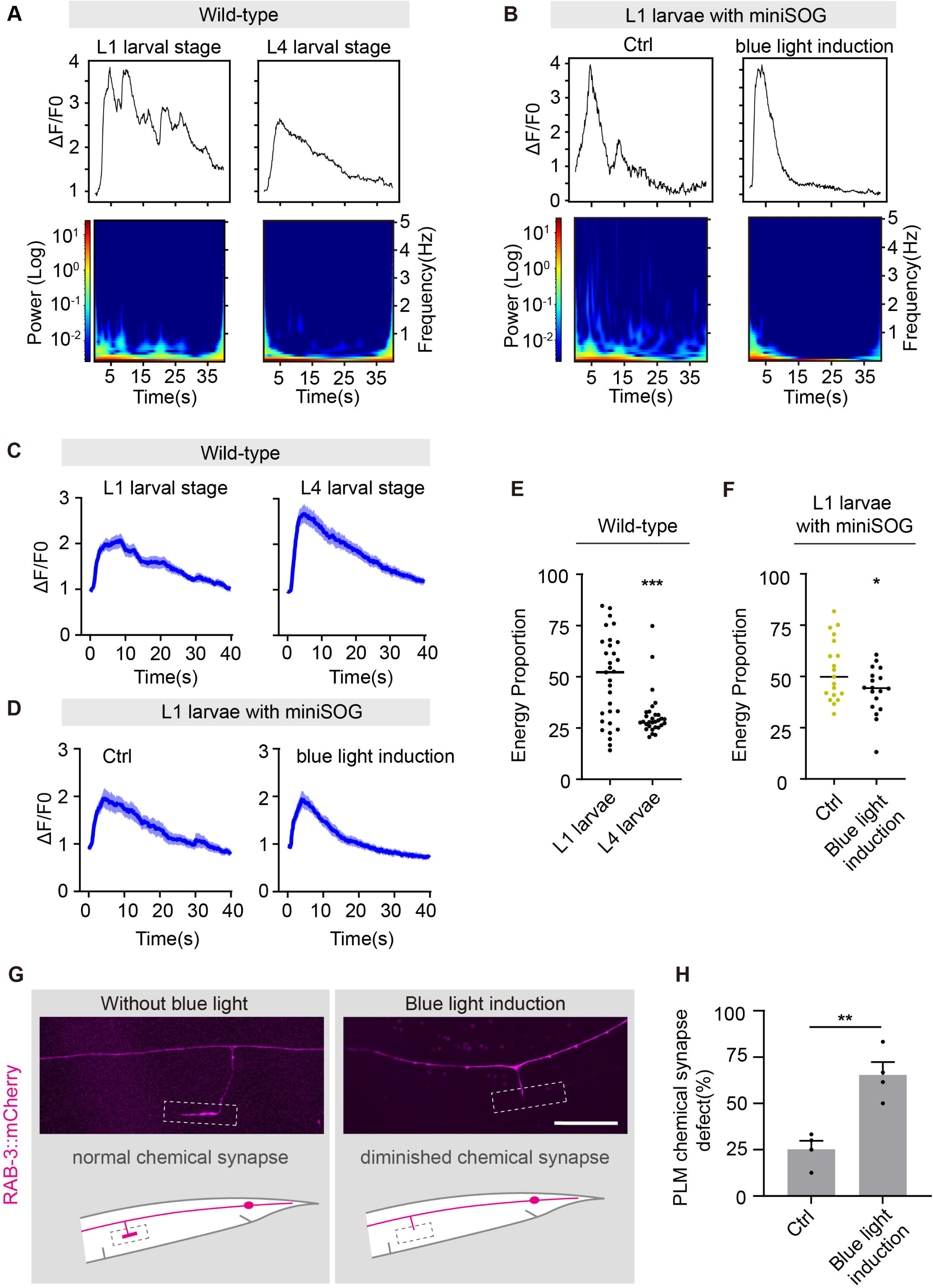
Transient electrical synapses coordinate calcium dynamics and regulate chemical synapse development. (A) Representative PLM calcium trace and wavelet spectrogram in L1 and L4 Gcamp6-expressing WT background animals. Calcium activity was quantified as changes in GCaMP fluorescence normalized to baseline (ΔF/F₀). (B) Representative PLM calcium trace and wavelet spectrogram in L1 *Pmec-4::miniSOG::unc-9 C. elegans* with or without blue light activation. (C) Population-averaged calcium trace of PLM neurons in L1 and L4 Gcamp6-expressing WT background animals. Blue line and shading represent mean ± SEM. n = 30 per stage. (D) Population-averaged PLM calcium trace of L1 *Pmec-4::mini-SOG* animals with or without blue light activation. n = 20 per group. (E) Quantification of secondary calcium signal energy proportion of PLM neuron in L1 and L4 Gcamp6-expressing WT background animals. The secondary calcium signal was defined as the ratio of summed energy of components >0.1 Hz during the secondary response window (5–30 s) to that during the full response period (2–38 s). n = 30 per stage. (F) Quantification of secondary calcium signal energy proportion of PLM neurons in L1 *Pmec-4::miniSOG::unc-9* animals with or without blue light activation. n =19 (without activation) and n = 18 (with activation). (G) Representative images of PLM chemical synaptic branch in L4 animals expressing miniSOG::UNC-9 and *mec-7* promoter-driven RAB-3::mCherry. The PLM chemical synapse was present without blue light induction but was absent after blue light induction. The schematic figure illustrates the normal PLM chemical synaptic and the synapse-diminished phenotype. The dashed box highlights the chemical synapse position. (H) Quantification of PLM chemical synapse defects in L4 animals expressing RAB-3::mCherry and miniSOG::UNC-9 with or without blue light induction. n = 38 (without induction) and n = 40 (with induction). For panels with quantification, error bars represent mean ± SEM. For confocal images, scale bar: 20 μm. Data in (E) and (F) were analyzed by Mann-Whitney U test. (H) was analyzed by Student’s t-test. *p < 0.05, **p < 0.01, ***p < 0.001.

We then assessed whether electrical synapses are required for these oscillations by performing optogenetic ablation using miniSOG-tagged UNC-9, and selectively disrupting L1 transient electrical synapses with blue light. While basal calcium signals remained unchanged (Fig. 2D), indicating that overall PLM neuron function was preserved after blue light treatment, wavelet analysis revealed that optogenetic ablation significantly reduced high-frequency calcium activity (Fig. 2B,F), confirming that transient electrical synapses mediate coordinated high-frequency calcium dynamics during early larval development.

To assess the biological significance of these electrical synapse-mediated calcium dynamics, we first examined neurite outgrowth and overall neurite length, but found no significant morphological defects (Fig. S2A,B,C,D). We next examined chemical synapse formation, given that electrical synapses regulate chemical synaptogenesis in many systems^31^ and PLM chemical synapses mature primarily during L1^32^. Analysis of chemical synapse formation using the presynaptic marker RAB-3 revealed striking defects. Control animals expressing miniSOG-tagged UNC-9 without blue light exposure displayed chemical synapse abnormalities in approximately 25% of PLM neurons (Fig. 2H). In contrast, optogenetic ablation dramatically increased this proportion to 65% (Fig. 2H), with many neurons exhibiting complete absence of RAB-3 puncta at expected synaptic locations (Fig. 2G). These findings indicate that transient electrical synapses during L1 are critical for proper chemical synapse formation, and their premature elimination disrupts the establishment of mature chemical connectivity.

### UNC-51 and UNC-76 function in the same genetic pathway to regulate electrical synapse elimination

To uncover the regulatory mechanism underlying PLM electrical synapse elimination, we performed an unbiased EMS (ethyl methanesulfonate)-induced forward genetic screening using GFP::UNC-9 transgenic line (*yadIs12*) worms to identify genes that suppress electrical synapse elimination. Through the screening, we isolated the mutant *yad29*, in which PLM neurons exhibited abnormally excessive electrical synapses throughout the entire neurite at L4 (Fig. 3A). Genetic mapping and rescue experiments identified *yad29* as a novel *unc-51* allele containing a single-nucleotide mutation introducing a premature stop codon at amino acid 672 (Fig. S3A). Consistently, two independent *unc-51* alleles, *unc-51(e1189)* and *unc-51(e369),* also exhibited similar excessive electrical synapses (Fig. 3A). Quantification in L4 revealed that 79.3%, 85%, and 87.4% of L4 animals exhibited the defect in *unc-51(e1189)*, *unc-51(yad29)*, and *unc-51(e369)* strains, respectively—all significantly higher than wild-type (Fig. 3B). Tissue-specific rescue experiments demonstrated that both pan-neuronal (*unc-33* promoter-driven) and PLM-specific (*mec-4* promoter-driven) UNC-51 expression significantly rescued the phenotype in *e369* worms, confirming a cell-autonomous requirement of UNC-51 for regulating PLM electrical synapse distribution (Fig. 3B).

**Figure 3.**
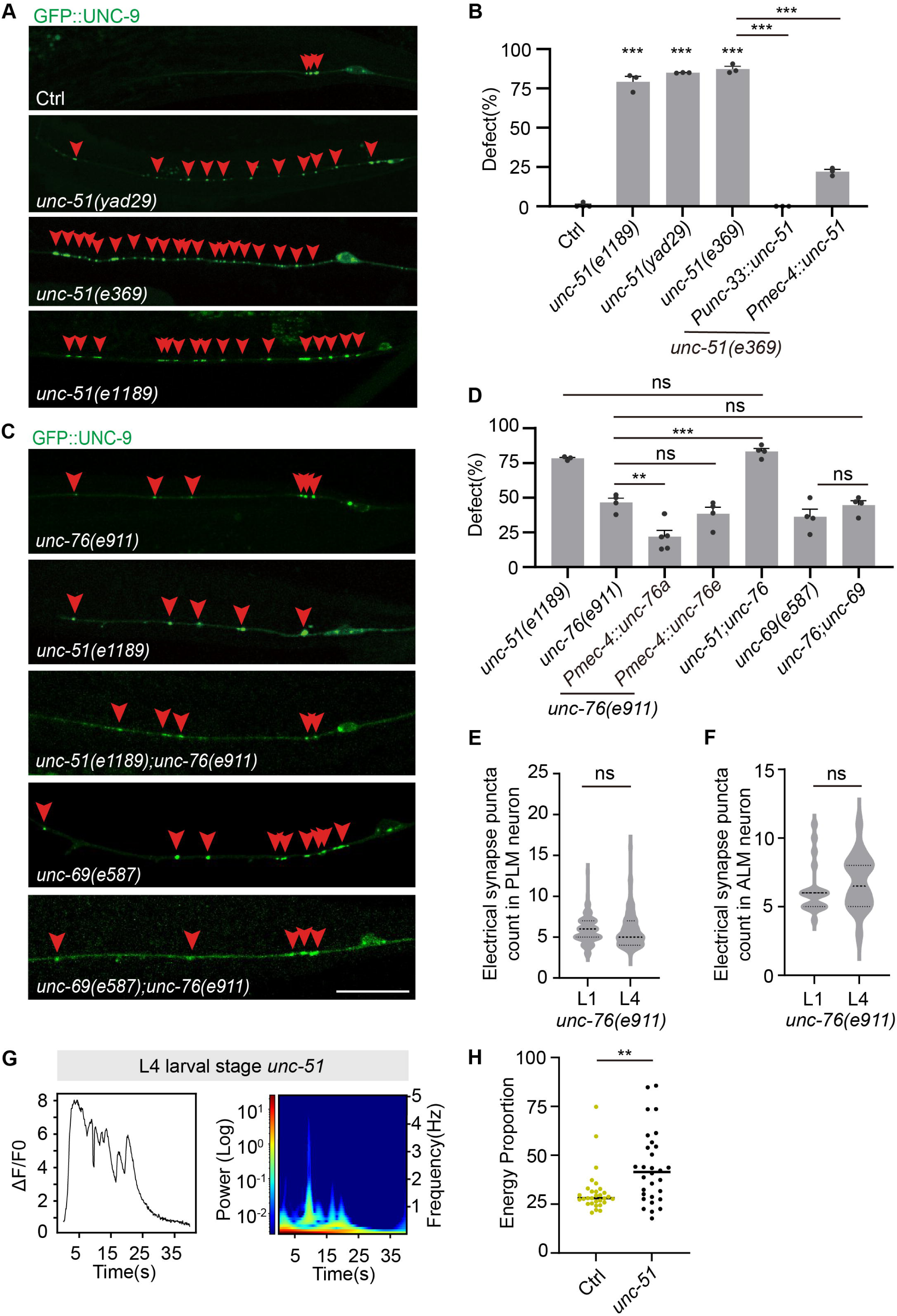
UNC-51 and UNC-76 regulate electrical synapse elimination in mechanosensory neurons. (A) Representative fluorescence images of GFP::UNC-9-labeled PLM electrical synapses in L4 control and *unc-51* mutant animals. Red arrowheads point to labeled electrical synapses. (B) Quantification of excessive PLM electrical synapse defects in L4 control, *unc-51* mutant, and *unc-51* rescue animals. n > 100 per genotype. (C) Representative fluorescence images of GFP::UNC-9-labeled PLM electrical synapses in L4 *unc-51*, *unc-76*, *unc-51; unc-76* double mutant, *unc-69*, and *unc-69; unc-76* double mutant animals. Red arrowheads point to labeled electrical synapses. (D) Quantification of excessive PLM electrical synapse defects in L4 *unc-51*, *unc-76*, *unc-76* rescue, *unc-51; unc-76* double mutant, *unc-69*, and *unc-69; unc-76* double mutant animals. n > 100 per genotype. (E, F) Quantification of GFP::UNC-9-labeled electrical synapses in PLM (E) and ALM (F) neurons of *unc-76* mutant animals at L1 and L4. For (E), n = 134 (L1), n = 105 (L4). For (F), n = 20 per stage. (G) Representative calcium trace and wavelet spectrogram of PLM neurons in L4 *unc-51* mutant animals. (H) Quantification of secondary calcium signal energy proportion in L4 control and *unc-51* mutant animals. n = 30 per genotype. For panels with quantification, error bars represent mean ± SEM. For confocal images, scale bar: 20 μm. Data in (B) and (D) were analyzed by one-way ANOVA with Tukey’s multiple comparisons test; (E) and (F) were analyzed by Student’s t-test; (H) was analyzed by Mann-Whitney U test. ns: not significant, **p < 0.01, ***p < 0.001.

UNC-51/ULK is an evolutionarily conserved kinase best characterized for its role in initiating autophagy^33^. To investigate how UNC-51 regulates electrical synapse development, we first accessed the involvement of autophagy by disrupting key autophagy-related genes (*atg-9*, *atg-6*, *atg-13*, *lgg-2*, *atg-4.1*). However, disruption of these autophagy genes did not reproduce the excessive electrical synapse phenotype (data not shown), demonstrating that UNC-51 regulates electrical synapses independently of its autophagy function.

Beyond autophagy, UNC-51/ULK has been implicated in vesicular trafficking and neuronal development^34–36^. To identify downstream effectors of UNC-51 in electrical synapse elimination, we systematically screened candidate genes implicated in regulating neuronal development. Among the genes tested, *unc-76(e911)* mutation produced excessive electrical synapse phenotypes resembling *unc-51* mutants (Fig. 3C). Quantification revealed that abnormal phenotypes were observed in 46.5% of *unc-76(e911)* mutants and 83.3% of *unc-76(e911); unc-51(e1189)* double mutants, a frequency not significantly different from the 78.3% observed in *unc-51(e1189)* mutants (Fig. 3C,D). These results indicate that UNC-51 and UNC-76 function within the same genetic pathway. Further screening of UNC-76-associated transport factors identified *unc-69* as an additional component of this pathway. *unc-69* encodes a coiled-coil protein homologous to mammalian SCOCO that directly interacts with UNC-76 to form kinesin-1 adaptor complexes mediating axonal transport^37,38^. In *unc-69(e587)* mutants, excessive electrical synapses were observed in 36% of animals, and double mutant analysis of *unc-76(e911)*;*unc-69(e587)* revealed 44.6% affected animals, which was comparable to single mutants (Fig. 3C,D), supporting that UNC-69 and UNC-76 function within the same regulatory pathway.

Collectively, these genetic analyses suggest that UNC-51 and UNC-76 act within the same genetic pathway governing electrical synapse distribution during neural development. To determine whether this pathway regulates developmental electrical synapse elimination, electrical synapse distribution across stages was examined in pathway mutants. Since *unc-51* mutants exhibit delayed mechanosensory neuron development at L1, we focused on *unc-76* mutants, which display relatively normal neurite development, to observe the elimination process. In both PLM and ALM neurons of *unc-76* mutants, the normal developmental reduction in electrical synapses was abolished (Fig. 3E,F, S3B), indicating that this pathway is required for developmental electrical synapse elimination.

To investigate the functional consequences of impaired electrical synapse elimination, calcium imaging was performed in PLM neurons of *unc-51* mutants. *unc-51(e1189)* mutants at L4 exhibited typical PLM calcium responses to mechanical stimulation, indicating that basic neuronal excitability was intact (Fig. S3C). However, wavelet analysis of spontaneous calcium activity revealed a significant difference: *unc-51(e1189)* mutant at L4 showed significantly elevated high-frequency calcium oscillations compared to wild-type controls (Fig. 3G,H), suggesting that persistent electrical synapses lead to abnormal neuronal activity.

### UNC-51 kinase controls electrical synapse elimination via UNC-76 phosphorylation

Having identified the genetic pathway, we next investigated the molecular mechanisms linking these components. To define the hierarchical relationship between *unc-51* and *unc-76*, the C-terminus of UNC-76 was endogenously tagged with mNeonGreen using CRISPR-mediated knock-in. The *unc-76::mNeonGreen* knock-in strain retained normal function, as evidenced by the absence of coiler phenotypes, with defect levels comparable to wild-type and significantly lower than those observed in *unc-76* mutants, in which 48.8% exhibited the defect (Fig. S4A). Using this functional reporter, UNC-76 protein expression levels were examined in different genetic mutant backgrounds to infer epistatic relationships. In *unc-51(e1189)* mutants, UNC-76::mNeonGreen levels were approximately five-fold higher than in wild-type neurons (Fig. 4A,B), demonstrating that UNC-51 negatively regulates UNC-76 protein abundance, and functions upstream of UNC-76 in the regulatory pathway.

**Figure 4.**
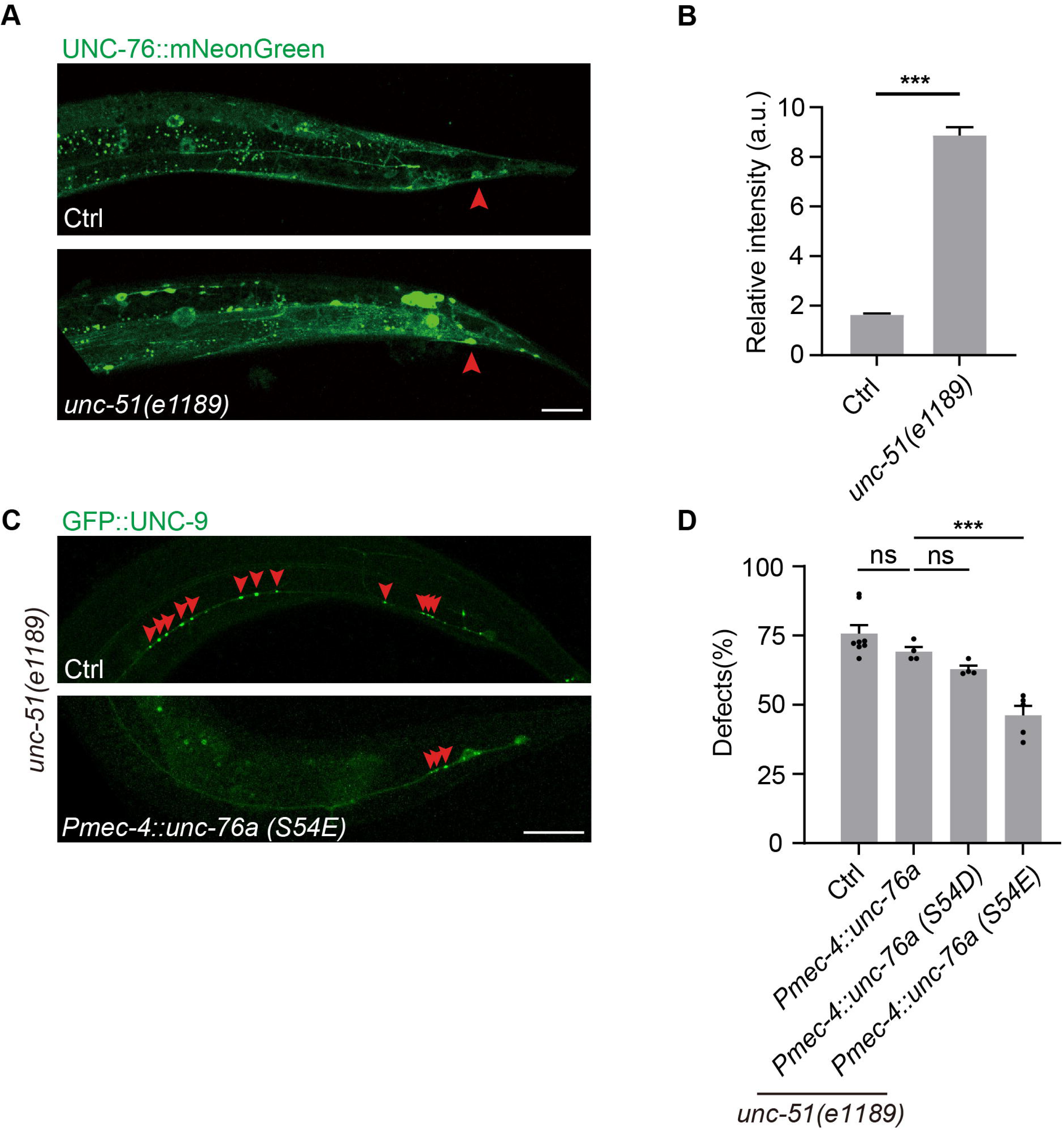
UNC-51-dependent phosphorylation regulates UNC-76 expression and function. (A, B) Representative fluorescence images (A) and quantification (B) of endogenous UNC-76::mNeonGreen in PLM neurons of L4 control and *unc-51* mutant animals. Red arrowheads in (A) indicate PLM neuron cell bodies. n = 103 (Ctrl), n = 105 (*unc-51*). (C) Representative fluorescence images of GFP::UNC-9-labeled PLM electrical synapses in L4 *unc-51* mutant animals and *unc-51* mutant animals expressing the UNC-76(S54E) phospho-mimetic mutants. Red arrowheads point to labeled electrical synapses. (D) Quantification of excessive PLM electrical synapse defects in *unc-51* mutant animals and *unc-51* worms expressing UNC-76 isoform a, UNC-76(S54D) phospho-mimetic, or UNC-76(S54E) phospho-mimetic mutants. n>100 per genotype. For panels with quantification, error bars represent mean ± SEM. For confocal images, scale bar: 20 μm. Data in (B) was analyzed by Student’s t-test; (D) was analyzed by one-way ANOVA with Tukey’ s multiple comparisons test. ns: not significant, ***p < 0.001.

Previous studies in *Drosophila* demonstrated that UNC-51 phosphorylates UNC-76 at Ser143 to regulate axonal transport by mediating motor-cargo assembly^34^. Notably, a conserved homologous phosphorylation site, Ser54, was identified in *C. elegans* UNC-76 long isoform (isoform a) (Fig. S4B, C). To determine whether UNC-51 regulates UNC-76 through phosphorylation, phosphomimetic mutants UNC-76(S54D) and UNC-76(S54E) were constructed, in which Ser54 was substituted with Asp or Glu to mimic constitutive phosphorylation. Remarkably, expression of UNC-76(S54E) in *unc-76(e911)* mutant worms partially rescued the excessive electrical synapse phenotypes, whereas wild-type UNC-76 expression showed no rescue effect (Fig. 4C,D). These results demonstrate that phosphorylation at Ser54 is both necessary and sufficient for UNC-76 function in electrical synapse elimination, suggesting that UNC-51 kinase directly phosphorylates UNC-76 to regulate electrical synapse elimination process.

### UNC-76 controls innexin transport and electrical synapse elimination

To determine the cellular basis of this phosphorylation-dependent regulation, we generated *Pmec-4::HaloTag::UNC-9; UNC-76::mNeonGreen* dual-labelled strains to simultaneously visualize both proteins. Time-lapse imaging of PLM axons revealed co-transport of UNC-9 and UNC-76 on moving vesicles in both anterograde (soma to axon) and retrograde (axon to soma) directions at L1 and L4 (Fig. 5A). Quantitative analysis revealed that transport patterns were consistent across developmental stages. In L1, anterograde co-transport occurred in 65% of cases, while L4 animals showed in 61% of cases, with no significant difference between stages (Fig. 5B). These findings demonstrate that UNC-76 directly participates in the bidirectional axonal transport of innexins.

**Figure 5.**
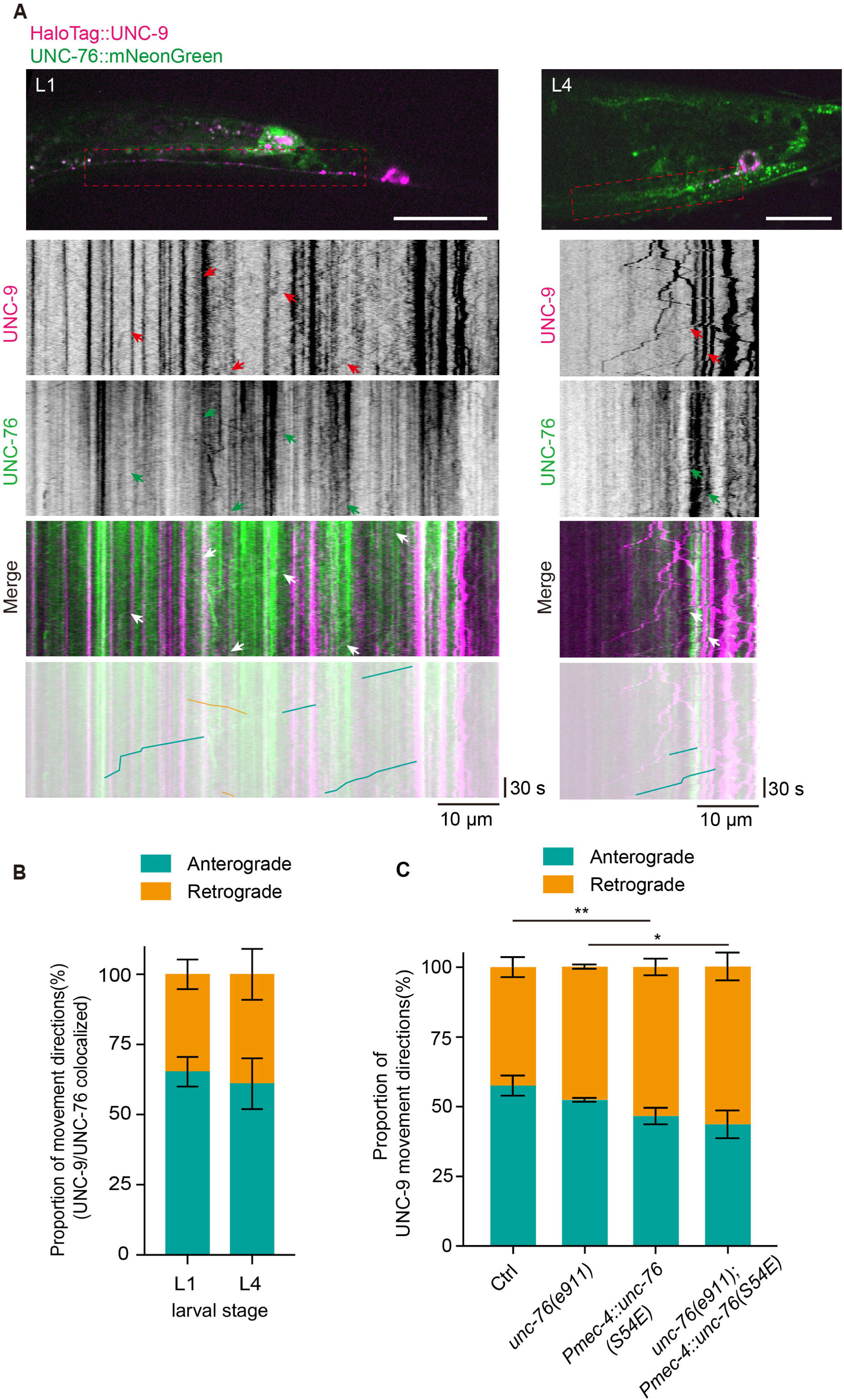
UNC-76 co-localizes with UNC-9 and exhibits stage-specific directional transport. (A) Representative fluorescence and time-lapse images of electrical synapse dynamics visualized using HaloTag::UNC-9 and endogenous UNC-76::mNeonGreen in L1 and L4 animals. The red rectangular box delineates the PLM axonal region used for time-lapse analysis. Arrowheads point to co-localized UNC-76 and UNC-9 puncta movement. The blue and yellow lines indicate UNC-9 anterograde and retrograde movement, respectively. (B) Quantification of the directional movement of co-localized UNC-76 and UNC-9 puncta in anterograde and retrograde directions at L1 and L4. n = 21 (L1), n = 20 (L4). (C) Quantification of anterograde and retrograde PLM UNC-9 movements in L1 control, *unc-76*, *pmec-4::unc-76(S54E),* and *unc-76; pmec-4::unc-76(S54E)* animals. n ≥ 17 per genotype. For panels with quantification, error bars represent mean ± SEM. For confocal images, scale bar: 20 μm. For time-lapse images, horizontal scale bar: 10 μm; vertical scale bar: 30 s. Data in (C) was analyzed by two-way ANOVA with Tukey’s multiple comparisons test. ns: not significant, *p < 0.05, **p < 0.01.

We next asked whether UNC-76 actively determines the directionality of innexin transport. We first assessed UNC-9 movements in *unc-76* loss-of-function mutants. We found that *unc-76* mutants displayed no significant alteration in the ratio of retrograde to anterograde UNC-9 movements compared to L1 controls (Fig. 5C), indicating that UNC-76 is dispensable for maintaining transport directionality at transient electrical synapses. We next expressed phosphomimetic UNC-76 (S54E) in both wild-type and *unc-76* mutant backgrounds. Expression of phosphomimetic UNC-76 significantly increased the proportion of retrograde UNC-9 movements both in wild-type animals and in *unc-76* mutants (Fig. 5C). The consistent gain-of-function effect across both backgrounds demonstrates that phosphorylated UNC-76 is sufficient to promote retrograde innexin transport. Together, these results indicate that UNC-76 participates in innexin transport but does not influence directionality in its unphosphorylated state. Phosphorylation of UNC-76 is sufficient to confer directional specificity, driving retrograde removal of innexins from electrical synapses and promoting their developmental elimination.

### UNC-76 associates with RAB-10-positive vesicles to mediate innexin removal

Having established that electrical synapse elimination requires active transport, the vesicular machinery responsible for this process was next investigated. Systematic screening of transport-related genes revealed that disruption of RAB-10 function by either *rab-10(dx2)* mutants or expression of GDP-locked, dominant-negative RAB-10(T23N) reproduced electrical synapse elimination defects (Fig. 6A,B). RAB-10 is a small GTPase known to regulate endosomal trafficking and has been implicated in receptor trafficking and degradation in both *C. elegans* and mammalian systems^39,40^. To determine whether RAB-10 functions in the same genetic pathway as UNC-51 and UNC-76, genetic interaction analysis was performed. Because *unc-51(e1189);rab-10(dx2)* and *unc-76(e911);rab-10(dx2)* double mutants exhibited severe sterility that precludes phenotypic analysis, we instead expressed a dominant-negative RAB-10(T23N) transgene in PLM neurons of *unc-51(e1189)* or *unc-76(e911)* mutant backgrounds as an alternative approach. Overexpression of RAB-10(T23N) in *unc-51* or *unc-76* mutants did not exacerbate the electrical synapse defects observed in the respective single mutants (Fig. 6A,B). This lack of additive effects suggests that RAB-10 functions within the same genetic pathway as UNC-51 and UNC-76.

**Figure 6.**
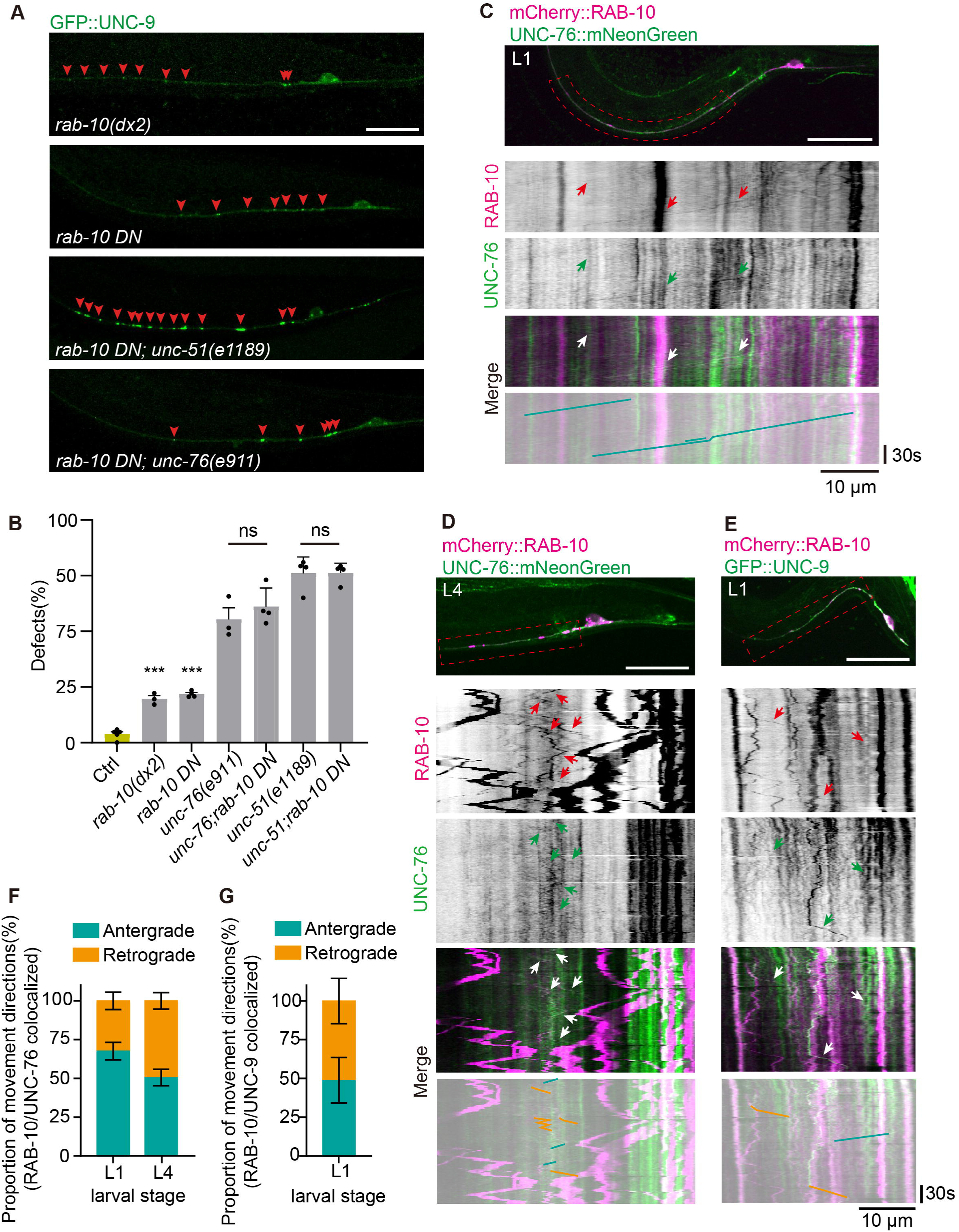
RAB-10 functions with UNC-51 and UNC-76 to regulate electrical synapse elimination. (A, B) Representative fluorescence images (A) and quantification (B) of GFP::UNC-9-labeled PLM electrical synapses in L4 *rab-10(dx2)*, *rab-10(DN)*, *rab-10(DN); unc-51(e1189)*, and *rab-10(DN); unc-76(e911)* animals. Red arrowheads in (A) point to labeled electrical synapses. n > 80 per genotype. (C, D) Representative fluorescence and time-lapse images showing co-localization of mCherry::RAB-10 and UNC-76::mNeonGreen puncta movement in PLM neurons at L1 (C) and L4 (D). Arrowheads point to co-localized puncta movements. The blue and yellow lines indicate anterograde and retrograde movement, respectively. (E) Representative fluorescence and time-lapse images showing co-localization of mCherry::RAB-10 and GFP::UNC-9 in PLM neurons at L1. (F) Quantification of the co-localized UNC-76 and RAB-10 puncta directional movement in anterograde and retrograde directions at L1 and L4. n = 16 (L1), n = 12 (L4). (G) Quantification of the co-localized RAB-10 and UNC-9 puncta directional movement in anterograde and retrograde directions at L1. n = 11. For all panels, error bars represent mean ± SEM. For confocal images, scale bar: 20 μm. For time-lapse images, horizontal scale bar: 10 μm; vertical scale bar: 30 s. Data in (B) were analyzed by one-way ANOVA with ’s multiple comparisons test; (F) were analyzed by Student’s t-test. ns: not significant, ***p < 0.001.

To determine whether RAB-10 works cooperatively with UNC-76, co-trafficking analysis using mCherry::RAB-10 in *unc-76::mNeonGreen* animals PLM neurons were performed. Time-lapse imaging at L1 and L4 revealed extensive co-trafficking of UNC-76 and RAB-10-positive vesicles in both anterograde and retrograde directions, with 68% anterograde movements in L1 and 51% in L4 (Fig. 6C,D,F), indicating that UNC-76 directly associates with RAB-10-positive vesicles in axonal transport. Additionally, co-localized axonal transport of RAB-10–positive vesicles with UNC-9 was also observed, with 49% of movements occurring in the anterograde direction (Fig. 6E,G), demonstrating a functional link between RAB-10-positive vesicles trafficking and innexin transport

Collectively, these findings establish RAB-10-positive vesicles as the key transport machinery that cooperates with UNC-76 to mediate active removal of innexin components during developmental electrical synapse elimination.

## Discussion

Widespread electrical coupling between developing neurons and their coordinated synchronous activity have been observed across systems, from the zebrafish spinal cord to the mammalian cortex^40,41^. However, how these transient connections are precisely terminated and the molecular mechanisms governing their elimination have remained unknown. Here, we identify the UNC-51/UNC-76 pathway as the core machinery for programmed electrical synapse elimination and demonstrate its functional necessity for proper circuit maturation (Fig. 7).

**Figure 7.**
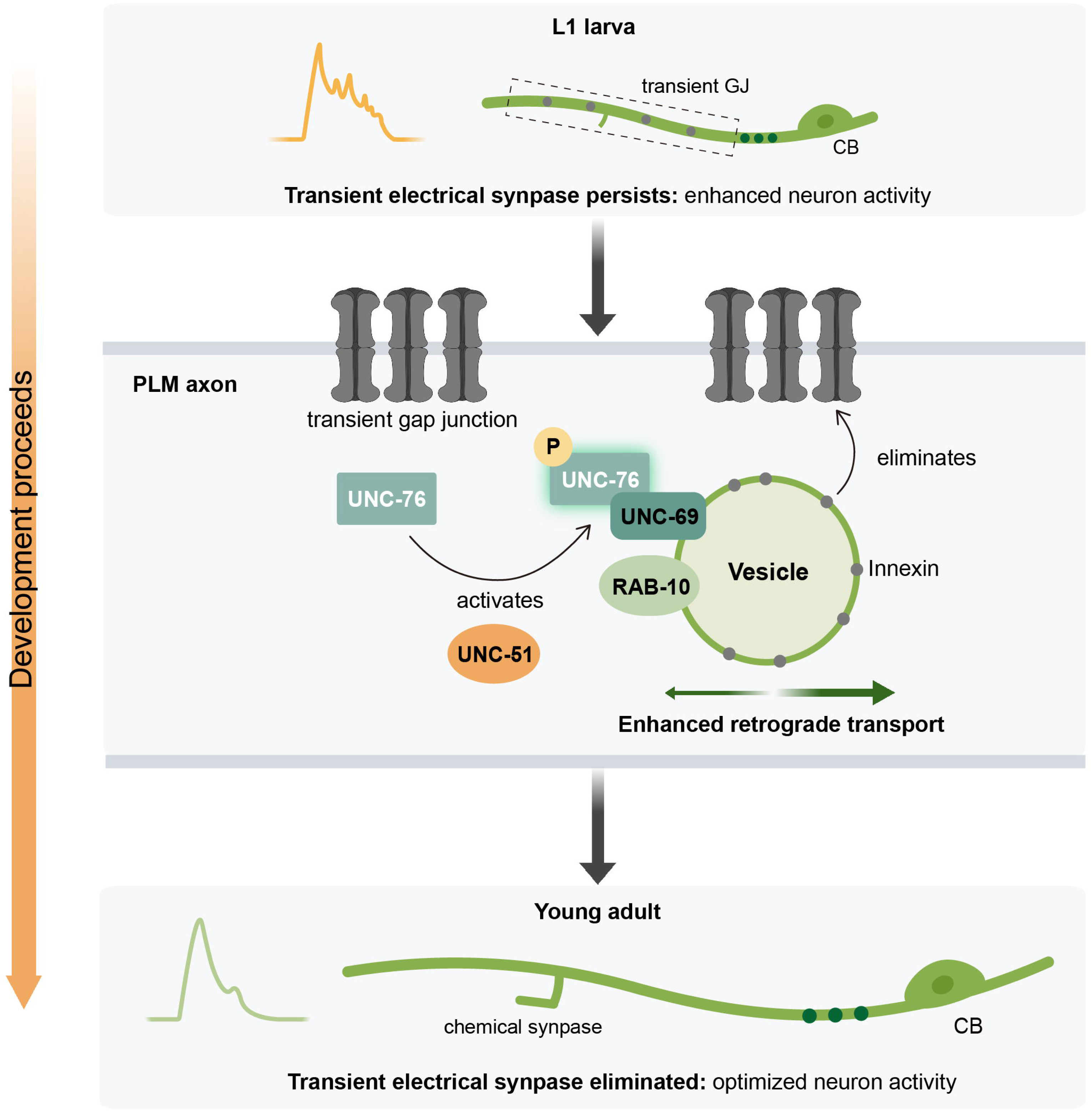
Schematic model illustrating the UNC-51–UNC-76–RAB-10 signaling pathway that regulates transient electrical synapse elimination, which is essential for normal neuronal activity and chemical synapse development.

### A phosphorylation switch converts constitutive trafficking into programmed elimination

Our findings reveal how developmental elimination is distinguished from constitutive turnover through temporal regulation of trafficking machinery. We propose that UNC-51-mediated phosphorylation of UNC-76 functions as a molecular switch that converts balanced bidirectional trafficking into directional removal (Fig. 7).

At early larval stages, electrical synapses in dynamic equilibrium are maintained by balanced trafficking. UNC-76 and RAB-10 continuously mediate both delivery and retrieval of innexins, resulting in stable synapse numbers despite rapid protein exchange. The regulatory transition occurs when UNC-51 kinase phosphorylates UNC-76 at Ser54, biasing the trafficking machinery toward retrograde transport and converting balanced flux into net protein removal. The economy of this mechanism is notable: the same molecular components execute both constitutive turnover and developmental elimination, with UNC-76 phosphorylation state determining the outcome. Retrieved innexins enter RAB-10-positive compartments, but whether these preferentially traffic to degradative versus recycling pathways during elimination is unclear. This model explains why disrupting any pathway component produces similar phenotypes and why rapid turnover coexists with stable synapse numbers at L1.

The architectural simplicity of electrical synapses—consisting primarily of clustered gap junction channels—makes cargo-selective transport particularly efficient. Unlike chemical synapse elimination, which requires multi-pathway mechanisms acting on diverse synaptic components, coordinated retrieval of the principal structural component is sufficient to dismantle the entire functional unit. For such architecturally streamlined synapses, cargo-selective vesicular trafficking thus offers an economical solution for developmental elimination.

### Temporal regulation of electrical synapse elimination is required for sequential circuit maturation

Transient electrical synapses in PLM neurons generate high-frequency calcium oscillations during early larval development, and their loss concurrently abolishes these dynamics and impairs chemical synapse formation. This provides the first evidence within a single genetically tractable system that electrical synapse loss simultaneously disrupts both processes, suggesting they may be coupled—though the precise causal relationship between calcium dynamics and chemical synaptogenesis remains to be established. Conversely, when elimination fails, persistent electrical synapses sustain calcium oscillations into later larval stages, causing neuronal hyperactivity—demonstrating that timely elimination is equally required to terminate early network activity and permit functional specialization. Together, these bidirectional perturbations reveal that proper circuit maturation depends not only on the early activity of transient electrical synapses, but on their precise developmental removal.

### Evolutionary conservation of the UNC-51/UNC-76 pathway

Our mechanistic analysis reveals that electrical synapse elimination requires active retrograde transport via the UNC-51→UNC-76→RAB-10 pathway. This pathway selectively removes innexins while presumably sparing other membrane components, indicating cargo specificity in the elimination machinery. UNC-51, the *C. elegans* homolog of mammalian ULK1/2, is best known for its role in autophagy initiation. However, our findings demonstrate an autophagy-independent function in regulating vesicular trafficking, consistent with emerging evidence that UNC-51/ULK proteins have diverse cellular roles beyond autophagy^34,41^. The net reduction in gap junctions suggests preferential removal, though the precise fate of retrieved innexins remains to be determined. The UNC-51 phosphorylation site on UNC-76/FEZ is evolutionarily conserved from invertebrates to mammals. Together with the established roles of ULK and FEZ proteins in mammalian axonal transport and synaptic development, this suggests that similar mechanisms may regulate connexin trafficking in vertebrate electrical synapses. Notably, connexin-based electrical synapses in mammalian brain also undergo developmental elimination in multiple regions, though the molecular pathways governing this process remain largely unknown. It will be important to determine whether ULK-mediated phosphorylation of FEZ proteins similarly regulates connexin trafficking in vertebrate neurons, which would indicate evolutionary conservation of this elimination mechanism from invertebrates to vertebrates.

### Broader implications for circuit refinement

Our work establishes cargo-selective retrograde transport as the mechanism for developmental electrical synapse elimination and provides the first molecular explanation for how transient synaptic architectures are precisely removed after completing their developmental functions. The finding that UNC-51 phosphorylation converts constitutive trafficking into active removal reveals a general principle: developmental transitions can be achieved by temporally regulating the activity state of constitutive cellular machinery. Given the evolutionary conservation of the pathway components and the widespread occurrence of electrical synapse elimination during mammalian brain development, these mechanisms likely represent fundamental principles of neural circuit assembly that extend beyond *C. elegans*.

## Materials and Methods

### *C. elegans* strains and genetics

*C. elegans* strains were maintained on NGM plates seeded with OP50 *Escherichia coli* at 20-23°C^42^. The N2 (Bristol) strain was used as the wild type. An unbiased genetic screen was performed using the NYL383(*yadIs12[Pmec-4::GFP::unc-9 + Pttx-3::RFP] IV)* strain following the standard ethyl methanesulfonate (EMS) mutagenesis protocol^43^. Mutant allele used in this study include *unc-51(e1189), unc-51(yad29), unc-51(e369), unc-76(e911), unc-69(e587), rab-10(dx2)*. A complete list of strains is described in Table S2.

### Plasmid construction and transgene generation

Plasmids were constructed using ClonExpress recombination technology (Vazyme, C112-01) or Gateway cloning technology (Invitrogen, 311791020). Gateway entry clones were generated using the backbone vector pCR8-JARID2 (Addgene, 114443), and destination vectors were derived from pCZGY553 (Addgene, 110883). Gene fragments were amplified from *C. elegans* N2 genomic DNA or cDNA.

For germline transformation, plasmids were microinjected at 10–50 ng/μL with co-injection markers (*Pttx-3::RFP*, *Prab-3::mCherry*, or *Podr-1::GFP*) at 10 ng/μL. Phosphomimetic UNC-76 variants (*Pmec-4::unc-76(S54D)* and *Pmec-4::unc-76(S54E)*) and dominant-negative RAB-10 (*Pmec-4::rab-10(T23N)*) were generated by site-directed mutagenesis of wild-type constructs using the Vazyme Mut Express II Fast Mutagenesis Kit (C214-01). All constructs were sequence-verified before microinjection. A complete list of plasmids is provided in Table S1.

### CRISPR/Cas9-mediated gene editing

CRISPR/Cas9-mediated genome editing was performed following the self-excising cassette (SEC) method described by Dickinson *et al*.^3^. Donor plasmid was constructed using the pDD282 backbone (Addgene, 66823), which contains the SEC. Homology arms (5′ and 3′) were amplified from N2 genomic DNA and cloned into pDD282 using Gibson assembly (Vazyme, C112-01). Plasmids expressing the Cas9 protein and single-guide RNAs (sgRNAs) were derived from pDD162 (Addgene, 47549) and generated using a site-directed mutagenesis kit (TOYOBO, SMK-101).

To generate the *unc-76*(caa97[*unc-76::mNeonGreen*]) knock-in strain, a microinjection mix containing donor plasmid (50 ng/µL; 852 bp 5′ homology arm + mNeonGreen + 669 bp 3′ homology arm), sgRNA-expressing plasmids (40 ng/µL; sgRNA#1: 5′-AATGTAGTCTGTCAGCAGGG-3′; sgRNA#2: 5′-TATTGTCATCGTGGATTGCG-3′), and Prab-3::mCherry co-injection marker (10 ng/µL; Addgene, 19359) was injected into young adult N2 hermaphrodites.

The *unc-9::47aa::mNeonGreen* and *unc-9::27aa::mNeonGreen* strains were generated using a similar strategy with sgRNA#1 (5′-CGTATGGTTGCAACTCACGC-3′) and sgRNA#2 (5′-GAGTTGCAACCATACGAAGT-3′), while *unc-9::4aa::mNeonGreen* was generated with sgRNA#1 (5′-ACGAGCTTGCGAAAACGTTA-3′) and sgRNA#2 (5′-AATTGAAAAAGCTACGAAGT-3′). Injected worms and their progeny were selected with hygromycin B (250 µg/mL) to identify roller individuals carrying successful integrations. Hygromycin-resistant rollers were screened by PCR genotyping, and the SEC cassette was removed by heat shock at 34°C for 6 h. Non-roller homozygous knock-in worms were confirmed by PCR genotyping. sgRNA sequences and CRISPR-related primers are listed in Table S3.

### RNA interference

RNA interference (RNAi) was performed by feeding worms with *E. coli* strain HT115 expressing double-stranded RNA (dsRNA). RNAi clones from the Ahringer feeding library^45^ were cultured overnight in LB broth containing ampicillin (100 µg/mL) at 37 °C. Bacterial cultures were concentrated and seeded onto NGM plates with ampicillin (100 µg/mL) and isopropyl 1-thio-β-D-galactopyranoside (IPTG, 5 mM), followed by overnight incubation at 37 °C to induce dsRNA expression. For neuron-specific RNAi, the transgenic strain *sid-1*(*pk3321*)*; uIs69[Punc-119::sid-1]* was used, in which RNAi sensitivity is restored exclusively in neurons^46,47^. *unc-22* RNAi was used as a positive control to assess RNAi efficacy. Worms were analyzed after being maintained on RNAi-expressing bacteria for at least two generations.

### HaloTag staining

For HaloTag::UNC-9 dynamic experiments, worms were incubated with 1 μM tetramethylrhodamine (TMR) HaloTag ligand (CA-TMR-1) in feeding culture at 20°C for 3 h in the dark. The feeding culture consisted of M9 buffer supplemented with centrifuged OP50 bacteria. Following incubation, worms were washed twice with M9 buffer and transferred to fresh NGM plates for 30 min at 20°C prior to imaging.

### Microscopy

Worms were immobilized on 6% agarose pads using either 1% 1-phenoxy-2-propanol (confocal imaging) or 2–8 mM levamisole (spinning-disk and widefield imaging) in M9 buffer. Confocal images were acquired using a Zeiss LSM880 microscope with a 40× oil-immersion objective. Time-lapse and optogenetic ablation-related images were acquired using a Nikon CWU-1 spinning-disk microscope equipped with a Hamamatsu ORCA-Fusion camera and a 100× oil-immersion objective (200 ms exposure per channel). Widefield calcium imaging was performed using a Zeiss Axioscope 5 microscope with a 63× oil-immersion objective (100 ms exposure, 20% LED illumination). Images were processed using ImageJ and Imaris Viewer.

### Calcium imaging and wavelet analysis

Imaging procedure followed established protocols^48,49^. GCaMP6s-expressing L4 hermaphrodites (n = 25) were immobilized on 5% agarose with 2 µM levamisole and mechanically stimulated by coverslip pressure. Changes were quantified as ΔF/F₀ with 5 s baseline periods. Cross-correlation analysis assessed PLM-PVC synchrony.

For wavelet-based signal analysis, the workflow was adapted from (Péter and Héja, 2024)^50^. ΔF/F₀ traces were calculated from raw fluorescence intensity values. Wavelet decomposition was performed using a complex Morlet wavelet (bandwidth = 1; center frequency = 1). The secondary calcium signal energy proportion was quantified as the ratio of the summed energy of frequency components above 0.1 Hz within the secondary response window (5–30 s) to that of frequency components above 0.1 Hz across the full response duration (2–38 s). Normality was assessed using the Shapiro–Wilk test. Since the dataset did not meet normal distribution criteria, non-parametric statistics were applied. Group comparisons were performed using the Mann-Whitney U test.

### Dendra2 photoconversion

For Dendra2 photoconversion experiments, pre-photoconversion images of PLM axons were acquired with both 488 nm and 561 nm channels (200 ms exposure). Photoconversion was performed by illuminating PLM axonal regions containing transient electrical synapses with 405 nm laser at 50% power for 20 s. Post-photoconversion imaging was immediately performed using identical settings. Worms were then transferred to NGM plates and maintained at 20°C. After 3 hours, PLM axons were re-imaged using the same parameters.

### miniSOG-mediated optogenetic ablation

For targeted miniSOG ablation, the strategy was adapted from previously described methods (Xu et al., 2013; Qi et al., 2012)^51,52^. Synchronized L1-stage *yadIS12* worms expressing *Pmec-4::miniSOG::unc-9* were mounted on agar pads and exposed to blue light to induce ablation. PLM axonal regions containing transient electrical synapses were selectively illuminated using the 488 nm laser line at 2 Hz (250 ms ON/ 250 ms OFF) with 80% laser power for 4 minutes, while avoiding adjacent regions to minimize off target photodamage. After illumination, animals were used for analyses or recovered from the agar pads and cultured under standard conditions.

### Image and quantitative analysis

For endogenous UNC-76 expression analysis in PLM cell bodies, the strain *unc-76(caa97[unc-76::mNeonGreen]*); *yadIs12[Pmec-7::mCherry; Pglr-1::mKate]* was used. The *unc-76::mNeonGreen* allele was generated by CRISPR/Cas9 knock-in, and the integrated *Pmec-7::mCherry* and *Pglr-1::mKate* transgenes facilitated PLM identification. Fluorescence images were acquired with 150 ms exposure and 80% LED illumination. The mNeonGreen fluorescence intensities were quantified using ImageJ. For each worm, three regions of interest (ROIs) were selected within the PLM cell body, and mean fluorescence intensity was calculated. Background fluorescence from non-neuronal regions was subtracted to obtain corrected intensity values.

For Dendra2::UNC-9 fluorescence quantification, fold-change analysis was performed at three time points: pre-photoactivation, 0 h post-photoactivation, and 3 h post-photoactivation. ROIs were defined within the PLM cell body and along photoconverted neurites, with ROI position and size kept constant across time points. Dendra2 fluorescence intensities from both green and red channels were measured at each time point. For each biological replicate, fold change was calculated as the fluorescence intensity at post-photoactivation time points relative to the pre-photoactivation value, which was set to 1 as the normalization baseline.

For calcium imaging fluorescence quantification, ROIs were defined within PLM cell bodies, and fluorescence intensity was acquired across time. Background fluorescence from non-neuronal regions was subtracted to obtain corrected values.

For PLM axon extension quantification, axon length was defined as the distance from the distal tip of the anterior axon to the vulva. Analyzed images were acquired with CWU-1 spinning-disk microscope 100× objective and axon length was measured using ImageJ.

### Statistical analysis

All data were analyzed using GraphPad Prism. Results are presented as the mean ± SEM. Statistical significance was determined using one-way analysis of variance (ANOVA) followed by Tukey’s post hoc test for multiple comparisons, or by two-tailed unpaired Student’s t-test and Mann–Whitney U test for pairwise comparisons. For all quantifications, P-values are indicated in the figures, where *P < 0.05, **P < 0.01, ***P < 0.001, and n.s. indicates not significant (P > 0.05).

---

**Figure S1.**
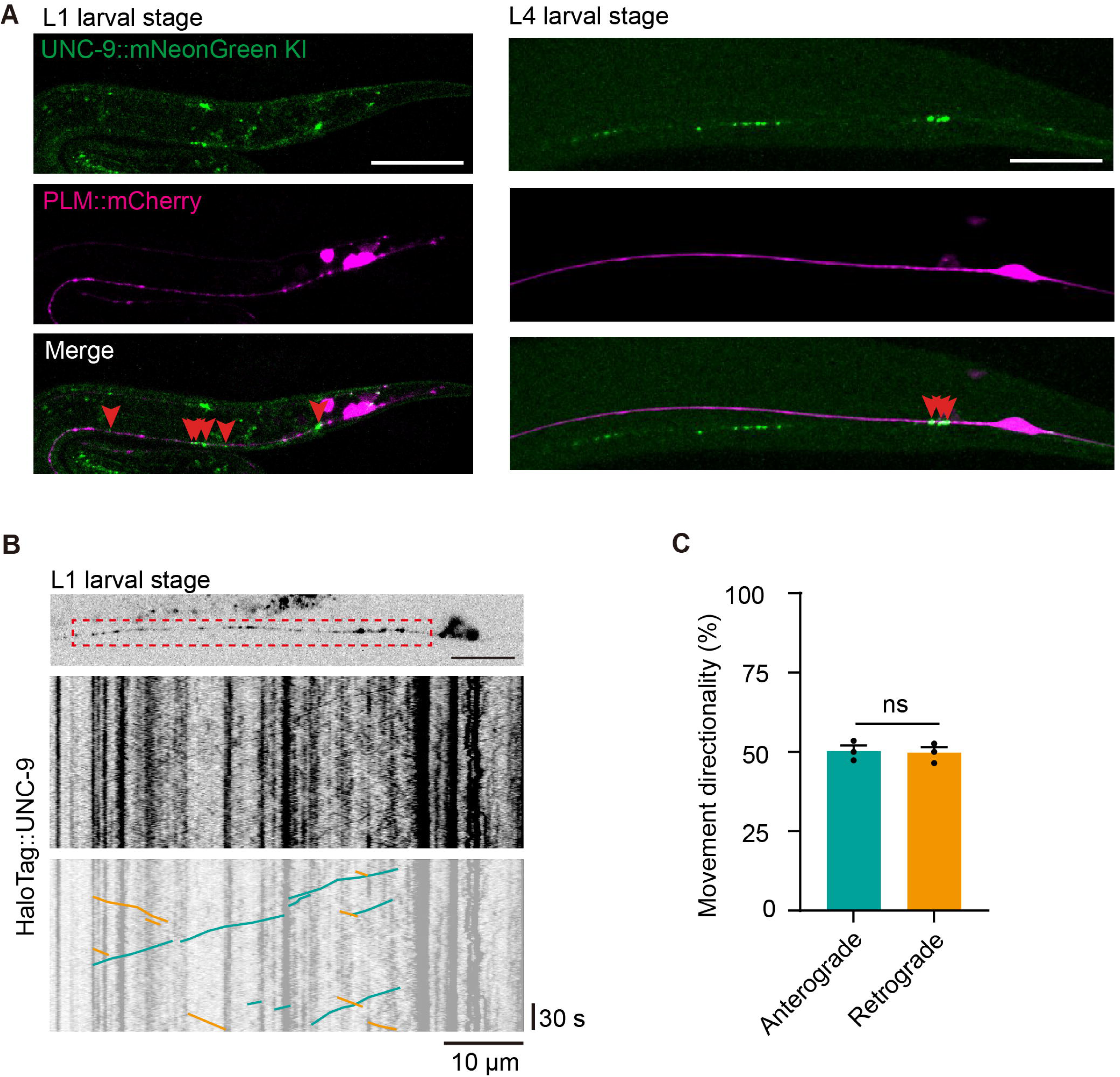
(A) Representative fluorescence images of endogenously C-terminally tagged UNC-9::mNeonGreen and *mec-4* promoter-driven mKate2 in PLM neurons at L1 and L4. (B) Representative time-lapse imaging of electrical synapse dynamics visualized using HaloTag::UNC-9 in WT animals. The blue and yellow lines indicate UNC-9 anterograde and retrograde movement, respectively. (C) Quantification of the directional movement of UNC-9 puncta originating from transient electrical synapses in L1 *pmec-4::Halotag::unc-9* animals PLM neurons. n = 17. For panels with quantification, error bars represent mean ± SEM. For confocal images, scale bar: 20 μm. For time-lapse images, horizontal scale bar: 10 μm, vertical scale bar: 30 s. Data in (C) was analyzed by Student’s t-test. ns: not significant.

**Figure S2.**
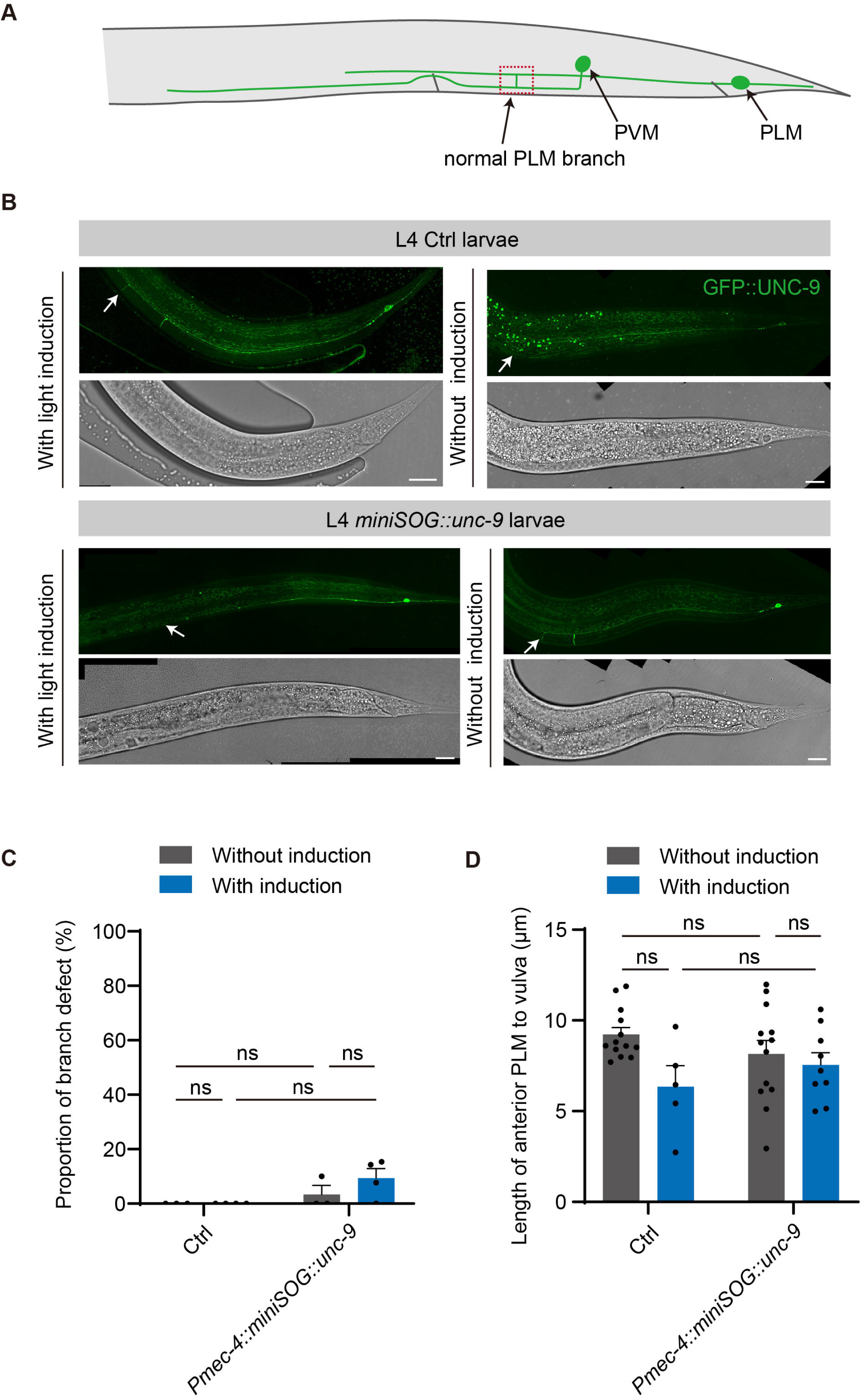
(A) Schematic figure of WT PLM neuron anterior axon morphology, showing a branch extending toward PVM and the axon terminus near the vulva. The dashed box highlights the PLM branch. (B) Representative fluorescence and DIC images showing PLM anterior axon branches in L4 Ctrl and *pmec-4:: miniSOG::unc-9* animals with or without blue light induction. White arrowheads point to axon branches. (C, D) Quantification of excessive axon branch defects (B) and the distance from the anterior PLM axon terminus to the vulva (C) in L4 Ctrl and *pmec-4:: miniSOG::unc-9* animals with or without blue light induction. For figure B, n > 23 per group. For figure C, n ≥ 5 per group. For panels with quantification, error bars represent mean ± SEM. For confocal images, scale bar: 20 μm. Data in (C) and (D) were analyzed by two-way ANOVA with the Tukey’s multiple comparisons test. ns: not significant.

**Figure S3.**
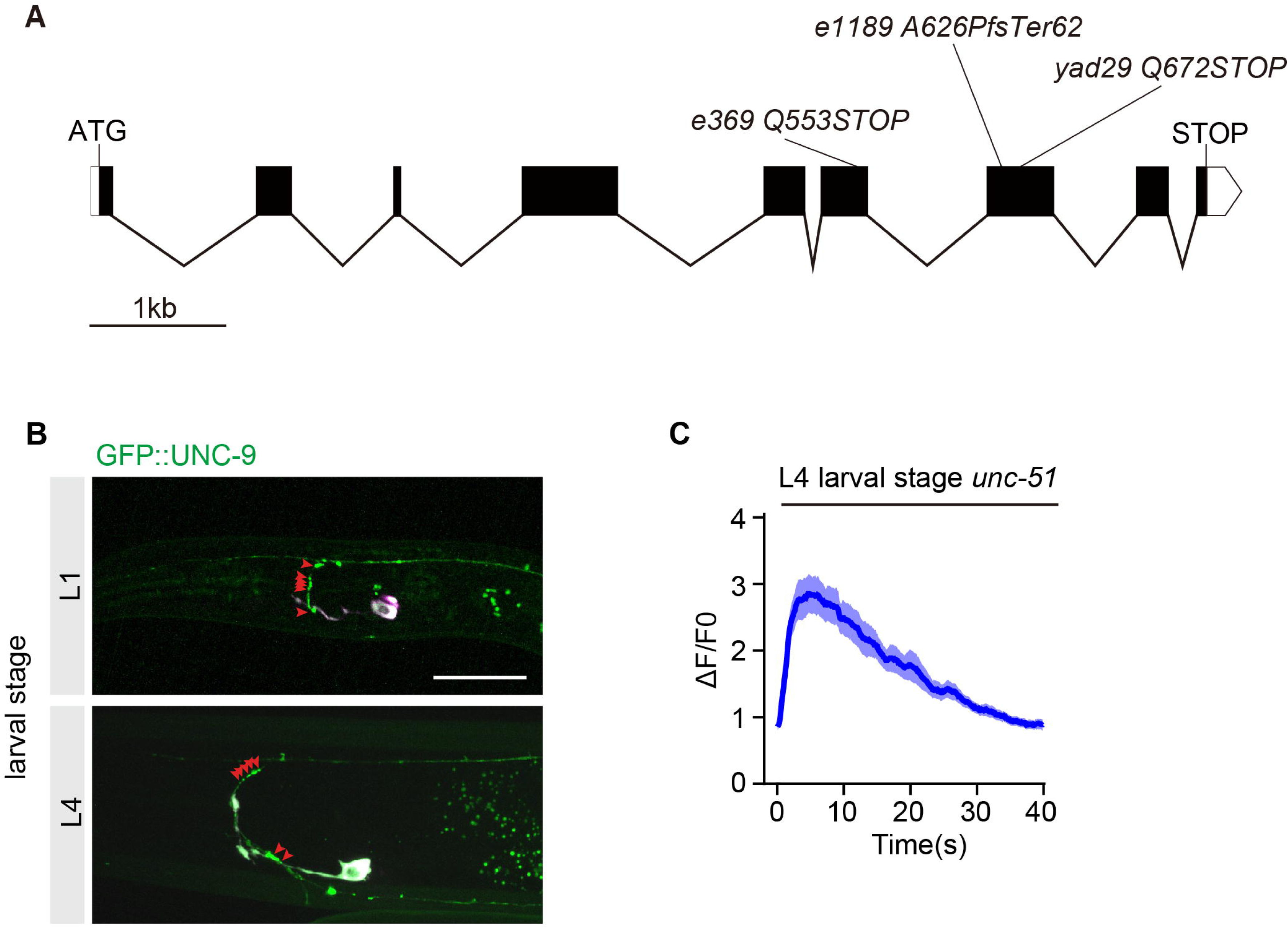
(A) Schematic diagram illustrating different *unc-51* alleles with their respective mutation sites and mutation types. Scale bar: 1 kb (B) Representative fluorescence images of GFP::UNC-9-labeled electrical synapses in ALM neurons at L1 and L4 in *unc-76* mutant animals. Scale bar: 20 μm (C) Population-averaged calcium trace of PLM neurons in L4 *unc-51* mutant animals. n = 30

**Figure S4.**
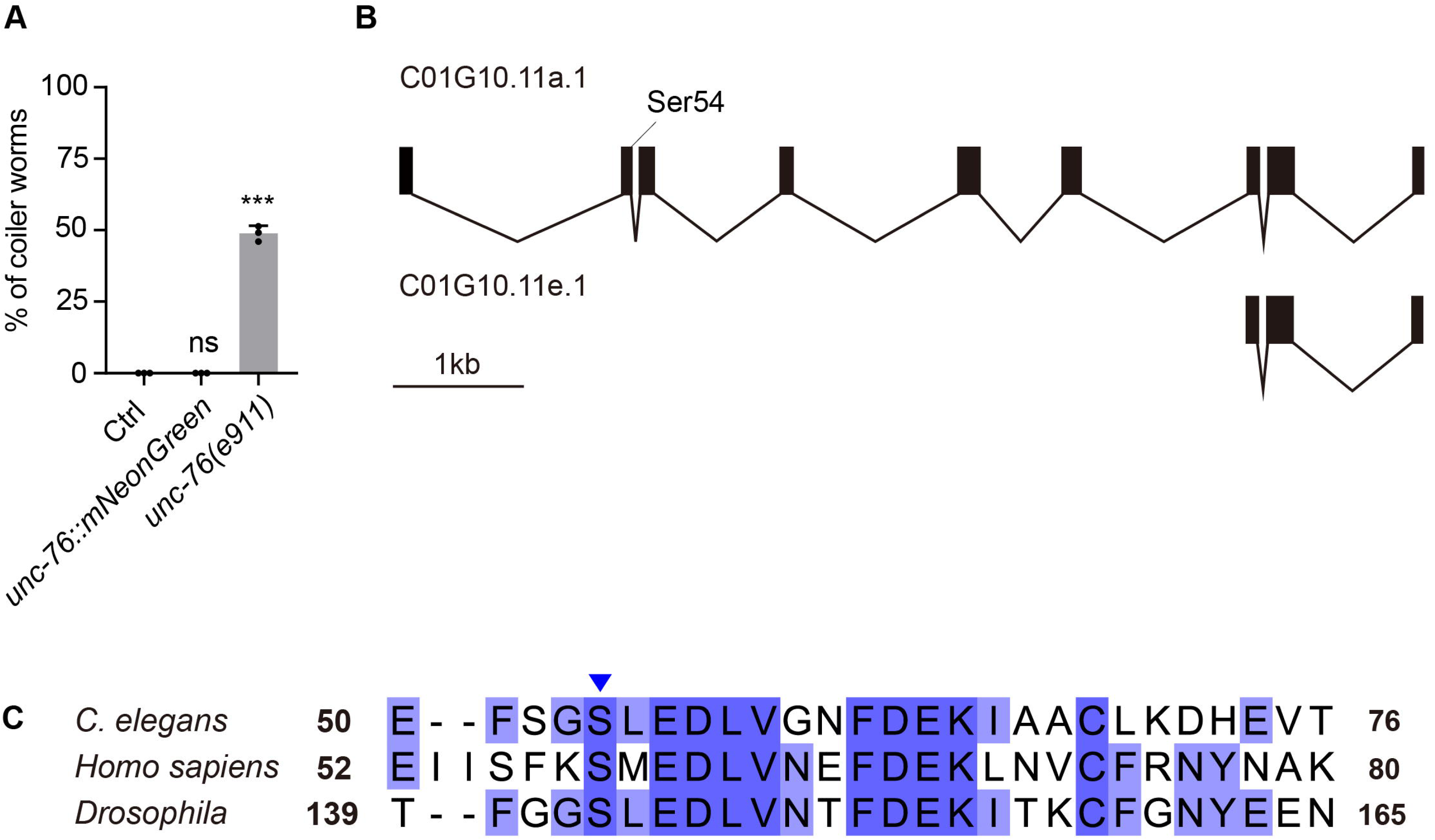
(A) Quantification of coiler locomotion defects in WT, endogenous *unc-76* C-terminal mNeonGreen knock-in, and *unc-76* mutant animals. n = 506 (WT), n = 449 (*unc-76::mNeonGreen*), n = 571 (*unc-76(e911)*). Error bars represent mean ± SEM. Data were analyzed by one-way ANOVA with Tukey’s multiple comparisons test. ns: not significant, ***p < 0.001. (B) Schematic diagram illustrating UNC-76 long isoform a and short isoform e, with the conserved serine 54 phosphorylation site indicated. Scale bar: 1kb (C) Sequence comparison of UNC-76/FEZ1 from *Drosophila*, *C. elegans*, and human, highlighting conserved regions surrounding the putative UNC-51 phosphorylation serine residue. The blue arrow point to the putative phosphorylation serine residue.

